# Mesoscale Modelling of Fibrin Clots: The Interplay between Rheology and Microstructure at the Gel Point

**DOI:** 10.1101/2024.09.20.614040

**Authors:** Elnaz Zohravi, Nicolas Moreno, Karl Hawkins, Daniel Curtis, Marco Ellero

**Affiliations:** Basque Center for Applied Mathematics (BCAM), Alameda de Mazarredo 14, Bilbao 48009, Spain; Medical School, Swansea University, Singleton Park, Swansea SA2 8PP, United Kingdom; Complex Fluids Research Group, Department of Chemical Engineering, Faculty of Science and Engineering, Swansea University, Swansea, SA1 8EN, United Kingdom; IKERBASQUE, Basque Foundation for Science, Calle de Maria Diaz de Haro 3, 48013, Bilbao, Spain

## Abstract

This study presents a numerical model for incipient fibrin-clot formation that captures characteristic rheological and microstructural features of the clot at the gel point. Using a mesoscale-clustering framework, we evaluate the effect of gel concentration or gel volume fraction and branching on the fractal dimension, the gel time, and the viscoelastic properties of the clots. We show that variations in the gel concentration of our model can reproduce the effect of thrombin in the formation of fibrin clots. In particular, the model reproduces the fractal dimension’s dependency on gel concentration and the trends in elasticity and gelation time with varying thrombin concentrations. This approach allows us to accurately recreate the gelation point of fibrin-thrombin gels, highlighting the intricate process of fibrin polymerization and gel network formation. This is critical for applications in the clinical and bioengineering fields where precise control over the gelation process is required.

## 1 Introduction

The viscoelastic properties of blood clots are biomarkers for blood coagulation in health and disease. They can reveal changes in clot structure, coagulation kinetics, and the processes of clot retraction and fibrinolysis.[1, 2, 3, 4] The mechanical properties of blood clots are determined by the microstructure of the complex fibrin network, which is formed by the polymerization of fibrinogen into fibrin fibers. Clots with altered fibrin microstructure have varying susceptibility to fibrinolysis. [5, 2, 3, 6] The fractal characteristics of incipient clots have been probed by oscillatory shear measurements of the gel point (GP) in whole blood,[7, 2, 8, 9] revealing the characteristic interplay between the structural properties and mechanical response. The clot’s mechanical response can change due to various conditions including venous thrombosis and coronary artery disease. In this context, the viscoelastic characterization of clots can be used in the screening and diagnosis of coagulopathies and in monitoring therapeutic interventions.

Fibrin-thrombin gels are a widely studied model system for blood clotting, providing insights into the clot’s microstructure and mechanical properties.[10, 8, 3, 11, 12] Thrombin plays a crucial role in the coagulation cascade by converting fibrinogen to fibrin. Experimental studies have shown that thrombin concentration can significantly alter the rate and microstructure formation of fibrin gels.[13, 10, 8, 14, 15] Investigations using Scanning electron microscopy (SEM) have shown that the concentration of thrombin significantly influences the characteristics of fibrin fibers, including their structure, length, and diameter. Specifically, lower thrombin concentrations result in thicker but less dense fibrin fibers, whereas higher concentrations lead to thinner, denser fibers. Similarly, the clots’ fractal dimension, *D*_*f*_, has been identified to correlate with thrombin levels.[16, 17, 18, 14]

Winter and Chambon[19] demonstrated that, at the gel point, both the storage modulus (*G*′) and the loss modulus (*G*″) exhibit a power-law dependence on the angular frequency (*ω*), with the relationship *G*″ ∼*G*′ ∼*ω*^*α*^, where *α* represents a stress relaxation exponent. This relationship indicates that, at the GP, the phase angle (*δ* = tan^−1^(*G*″*/G*′)) becomes independent of frequency and equals *δ* = *απ/*2. Evans et al [20] demonstrated that the GP of coagulating blood could provide a robust measure of a ‘clotting time’ and that power-law exponent (*α*) is sensitive to variations in thrombin availability. This sensitivity suggests that *α* can serve as a critical parameter for investigating and monitoring the impact of thrombin on the microstructure and stability of clots in a clinical setting.[2, 20]

Over the last decade, different numerical and experimental investigations have provided insights into blood clot rheological and microstructural characteristics. [18, 12, 14, 21] In the context of fibrin clots at the gel point, relevant numerical studies have been also addressed. Curtis and coauthors[3, 11] have explored the formation of incipient clots in fibrin-thrombin gels and heparinized blood using molecular dynamics simulations. They observed a correlation between gel-point time (tGP) and fractal dimension, noting that *D*_*f*_ decreases as tGP increases. Although their numerical model provides valuable insights into the microstructural properties of these gels, it does not account for variations of the viscoelastic properties with changing thrombin concentration. Overall, there is a noticeable knowledge gap in the numerical modeling of fibrin-clot formation. Existing models fail to simultaneously account for changes in the gel structure and viscoelastic response as thrombin concentration varies. Moreover, they do not adequately capture the key mechanical and structural relationships observed at the gel point. Here in, we provide a numerical scheme that captures relevant rheological and structural properties of fibrin-thrombin network formation. We demonstrate that the experimentally reported effects of thrombin concentration on the fractal dimension, gel elasticity, and gel time of fibrin-thrombin gels are accurately reproduced.[3, 11]

## 2 Definition of Gelation System

Gel-network formation can be modeled using explicit coarse-grained representations of the aggregating molecules. These models are effective for capturing the specific interactions and kinetics of molecule aggregation,[22, 23] as well as the distinct morphological features of these molecules.[24, 25] However, the computational demands of such detailed simulations can limit their applicability to larger spatial and temporal scales. Here, we adopt a recently proposed mesoscale framework[26] to model biological clustering, accounting for hydrodynamic interactions and network formation kinetics. This approach balance detailed microstructural features and computational efficiency, enabling the study of morphology and rheological response of the gel over larger scales. The framework discretizes the gelling system into a particle-based representation, in which the microstructural elements are constituted by passive (**P**), active (**A**), and solvent (**S**) particles. The particle interactions are modeled using the Smoothed Dissipative Particle Dynamics method[27], and the connectivity (bonding) between particles uses Morse potentials[28] (see simulation method for a detailed description of the model). As described by Zohravi et al.,[26] the rate of bond formation and maximum coordination number of bonds can be adjusted to model a variety of biological clustering mechanisms and morphologies.

The formation of the fibrin gel is a hierarchical process that involves the aggregation of fibrin monomers into mesoscale-mesh structures, which subsequently polymerize into fibers and crosslink to form a network of fibers that constitute the clot, as illustrated in Fig. 1.a. Here, **P** particles act as mesoscale-mesh subunits that polymerize into fibers by attaching to **A** particles. These active particles, once part of the fibers, can then cross-link to form the final gel structure (see Fig. 1.a).

**Figure 1:**
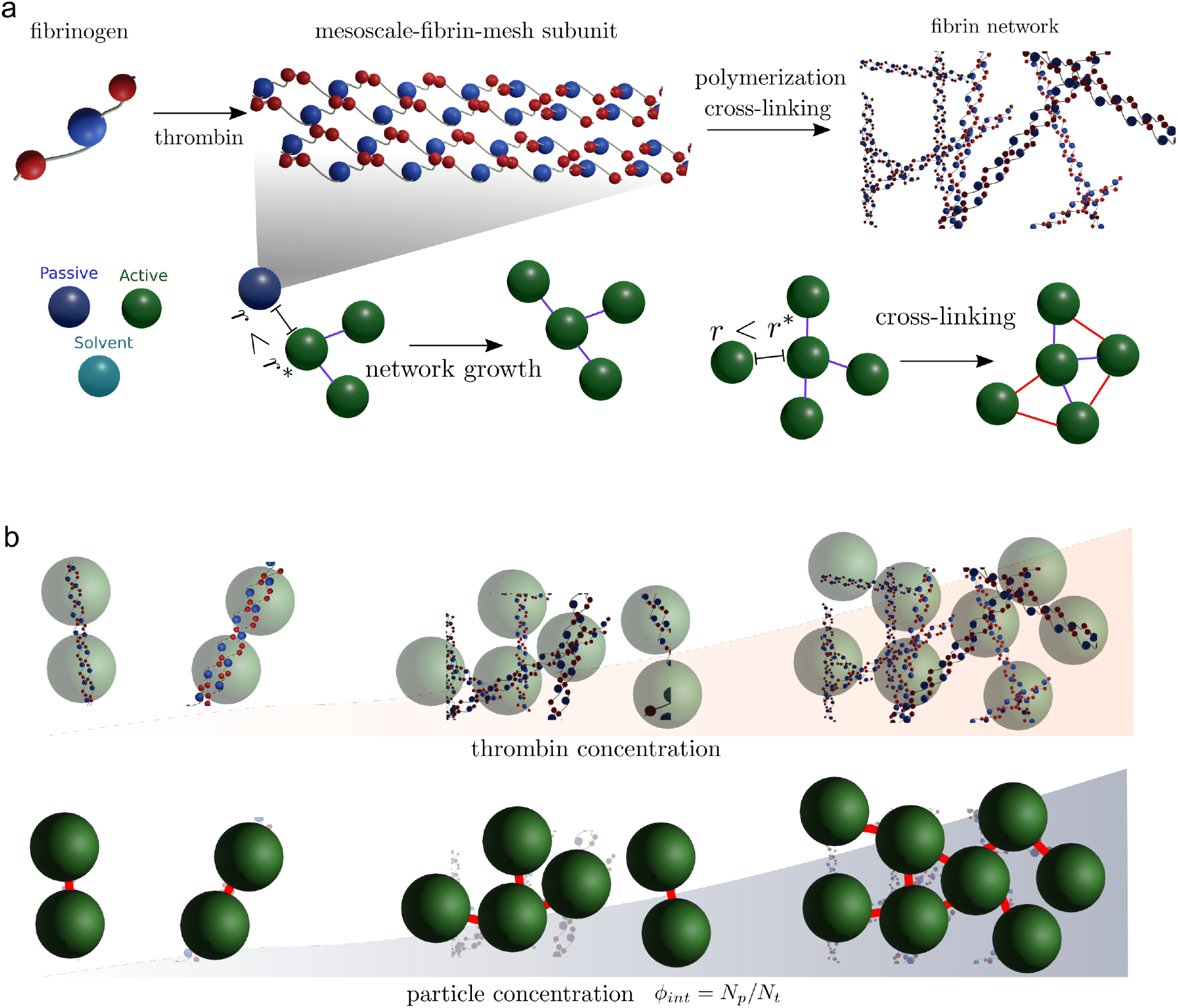
Schematic of the fibrin-gel formation process: (a) Hierarchical aggregation of fibrin monomers into mesoscale-mesh structures, followed by polymerization into fibers and crosslinking to form a network. Passive particles (**P**) polymerize into fibers upon bonding with active particles (**A**), which then cross-link to finalize the gel structure. (b) The initial concentration of passive particles (*ϕ*_int_) serves as a proxy for thrombin concentration. A higher *ϕ*_int_ reflects an increased thrombin concentration, due to the larger probability to form mesoscale-mesh subunits.

Therefore, we can describe the fibrin-gel formation in two main stages: *activation*, where passive particles become active by bonding with active particles, and *cross-linking*, where these now-active particles bond among themselves to solidify the structure. Initially, only passive and solvent particles constitute the simulated system. The gelation initiates by placing a single active particle seed in the center of the domain.

Previous studies in fibrin-thrombin gels have shown that the fractal dimension, elasticity, and gelation time are sensitive to the activation and crosslinking mechanisms.[3, 11, 29]. Based on the framework introduced by Zohravi et al.,[26] we investigate three representative clustering mechanisms, to identify the model that better captures the viscoelastic transitions during the gel formation dynamics. These mechanisms are denoted as *Branched, Low-Connected*, and *Highly-Connected*, and allow us to sample different degrees of connectivity in the system and cross-linking rates.

During the activation stage, the bonding between passive and active particles occurs when particles are within a distance *r*^∗^ of each other. The bond formation process, repeats with nearby active particles, allowing for a maximum of three bonds per particle. During the cross-linking stage, the bonding between **A** particles also takes place when their distance is smaller than *r*^∗^. Depending on the clustering mechanism a maximum number of ”*m*” bonds can be formed per **A** particle. We set *m* = 0 for branched, *m* = 2 for Low-Connected, and *m* = 10 for Highly-Connected. This allows us to mimic the various degrees of connectivity between fibers.[30]

It is important to note that although activation and cross-linking can occur synchronously, they are consecutive stages from the **P** particle standpoint. As described by Zohravi et al.,[26] is it possible to control the rate of the different stages, using a lag time between activation and crosslinking, or equivalently changing the bonding probability of each stage. If the lag time is too short, the crosslinking stage can overtake the gelling process, leading to smaller aggregates that do not percolate the domain; conversely, if the lag time is too long, the formed gels get kinetically trapped in the branched condition. Here, we set a bonding probability of one for the activation stage –as long as the particles are within the bonding distance, they will form a bond-. Whereas, the bonding probability is reduced to 1*/*100 for cross-linking to mimic a slower cross-linking rate than activation.

We denote the initial concentration of **P** particles as *ϕ*_int_ = (*N*_*p*_*/N*_*t*_)*×*100%, where *N*_*p*_ and *N*_*t*_ are the numbers of passive particles and total particles in the system, respectively. Here, we use *ϕ*_int_ as a proxy for thrombin concentration, and we investigate the effect of varying *ϕ*_int_ on the gelation process, as illustrated in Fig. 1.b. Thrombin is responsible for converting fibrinogen to fibrin, consequently generating the mesoscale-fibrin-mesh subunits. Therefore, it is expected that increasing thrombin concentration correlates with a higher number of initial passive particles. Overall, increasing *ϕ*_int_ implies that the probability of activation of passive particles increases. Similar approaches have been successfully used to model the effect of thrombin on the formation of fibrin-thrombin gels.[3, 11] In the Results section, we show that *ϕ*_int_ effectively captures the structural and rheological effects of thrombin on the gelation process.

## 3 Simulation Method

We use the smoothed dissipative particle dynamics (SDPD)[27, 31] method to model the hydrodynamic interaction between the particles that constitute the system. SDPD is a thermodynamically consistent method that discretizes the fluctuating Navier-Stokes equations and has been widely used in the modeling of soft matter.[32, 33, 34] We consider a gelling system containing *N* particles with a volume *V*_*i*_, such that 1/*V*_*i*_ = *d*_*i*_ = Σ_*i*_ *W* (*r*_*ij*_, *h*), being *d*_*i*_ the number density of particles, *r*_*ij*_ = |**r**_**i**_ − **r**_**j**_|, and *W* (*r*_*ij*_, *h*) an interpolant kernel with finite support *h* and normalized to one. The evolution equations for the position of the particles is d**r**_*i*_*/*d*t* = **v**_*i*_, whereas the stochastic differential equation of the momentum, can be expressed as

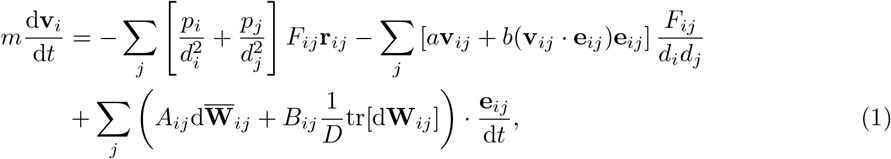

where **v**_*i*_ and *p*_*i*_ are velocity and pressure of the *i*-th particle. **v**_*ij*_ = **v**_*i*_ − **v**_*j*_, **e**_*ij*_ = **r**_*ij*_*/*|**r**_*ij*_|, *a* and *b* are friction coefficients related to the shear *η* and bulk *ζ* viscosities of the fluid through *a* = (*D* + 2)*η/D* − *ζ* and *b* = (*D* + 2)(*ζ* + *η/D*). *D* is the dimension of the system. In Eq.(1), we conveniently introduce the positive function *F*_*ij*_ = −∇*W* (*r*_*ij*_, *h*)*/r*_*ij*_. The last term in Eq.(1), consistently incorporates thermal fluctuations in the momentum balance. SI.S1, Eqs.(S1-S5) provide a detailed explanation of the terms *A*_*ij*_ and *B*_*ij*_, the equation of state that defines the pressure *p*, and the form of the function *F*_*ij*_.

The bonding between connected particles is modeled using a Morse potential[28] of the form 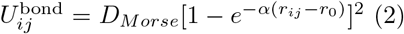. The Morse potential is an ideal and typical anharmonic potential among the several molecular potentials, resulting in a bonding force 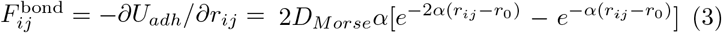. Parameters used in this potential are as follow: bond strength, *D*_*Morse*_ = 300, stiffness parameter, *α* = 1, equilibrium bond distance, *r*_0_ = 0.25 and bond criteria, *r*^∗^ = 0.4. Additional information about implementing the SDPD model is produced in SI.S3.

## 4 Gel Characterization

The gelling mechanisms adopted allow us to model gels with different rheological responses and microstructural connectivity. To investigate the rheological response of the gels, we conduct the small-amplitude oscillatory shear (SAOS) analysis, as described in the following section. Whereas to account for variations in the microstructural features of the gels, we describe the connectivity in terms of their bond distribution, and the characteristic average number of bonds between particles-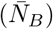 and maximum-number (*N*_*B*−*Max*_) of bonds per particle. In addition to the bond distribution, we also estimate the fractal dimension of the different gels, as an indicator of morphological distribution of the network. See SI.S2 for a detailed description of the fractal dimension computation. Given the statistical variability of the gels (and the stochastic nature of SDPD) depending on the initial distribution of the particles, the different gel characteristics are determined as mean values of five independent realizations of the gelling process.

### 4.1 SAOS Analysis

We use SAOS[35, 36] to study the linear viscoelastic response (complex modulus and loss tangent) for our model gels. We define the gel time (tGP) as the time at which the sol-gel transition occurs due to network formation. The selection of the waveforms’ upper (5*π* Hz) frequency limit is determined by considering the effects of fluid inertia. The lower limit, (0.25*π* Hz) is selected for computational reasons. Lower frequencies pose numerical challenges, due to excessively long simulation times, or very small changes in the system behavior, making it difficult for our numerical method to detect meaningful responses. We first determine the ‘linear viscoelastic region’ for our gels by performing amplitude sweeps at a fixed frequency. In the linear viscoelastic limit, the moduli do not depend on the amplitude of oscillation. A strain amplitude of 0.05 was found to be well within the linear-viscoelastic response throughout the gelation process. To ensure a quasi-incompressible flow, the speed of sound was set to *c* = 50 ≫ *V*_*max*_(*ω*) = *γ*_0_*ω* for every frequency, such that the Mach number was always smaller than one. In addition, the solvent viscosity considered produces a Reynolds number *Re* = *ρLV*_*max*_*/η* always smaller than one for all the frequencies investigated.[37] See SI.S4-S5 for a detailed description of the simulation setup and stress measurement. To streamline the presentation of our results, we adopt non-dimensional time units, by normalizing the time with the corresponding SDPD viscous time, *t*_*v*_ = (*h*)^2^*ρ/η*, where *h* is the kernel cutoff radius, *ρ* and *η* are the density and viscosity of the solvent, respectively. To account for the difference in bonding rate between activation and cross-linking, the lag time between the two stages corresponds to 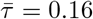. We normalize the elastic modulus, viscous modulus, and frequency by utilizing the values Böhme and Stenger proposed in their work[38]. 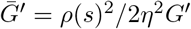 and 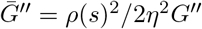 are the dimensionless elastic and viscous modulus. The dimensionless frequency is 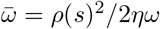 and *s* is the gap between the fixed plate and the moving plate in the SAOS test. We conduct simulations for gel concentrations of *ϕ*_*int*_ = 7.5, 10, 12.5, 15%. Further details related to the maximum value of gel concentration can be found in Section SI.S2.

## 5 Results

We conduct an initial assessment of the connectivity and viscoelastic properties of the different gelation mechanisms (Branched, Low-Connected, and Highly-Connected) to identify which model better captures the dynamics of fibrin-gel formation. In Fig. 2.(a), we present the variation in the connectivity features for the three mechanisms at their final state. We define the final gel state as the point where bond formation stabilizes. Specifically, we consider the gel to be in its final state when the number of new bonds changes by less than 0.1% over 10000 time steps. Their characteristic average bonds between particles and maximum number of bonds per particle can be summarized as follows:

**Figure 2:**
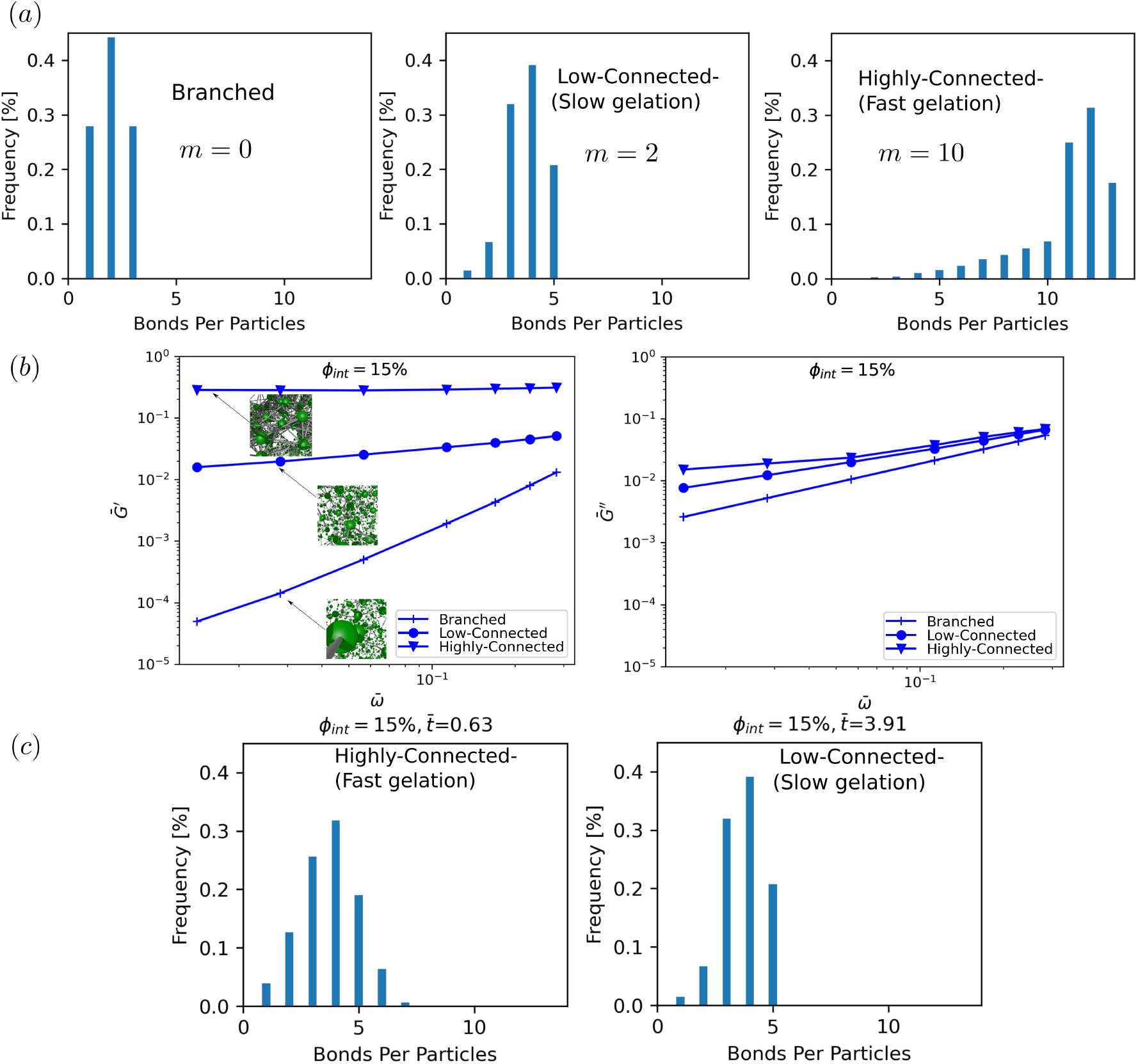
(a) Bond histogram of 3 proposed mechanisms. (b) Elastic and viscous modulus of 3 proposed mechanisms at *ϕ*_*int*_ = 15% and their corresponding gel morphology. (c) Comparing the bond histogram for the Highly-Connected mechanism and the Low-Connected mechanism at the same average number of bonds 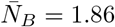

Branched: 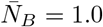, *N*_*B*−*Max*_ = 3

Low-Connected: 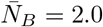, *N*_*B*−*Max*_ = 5

Highly-Connected: 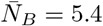, *N*_*B*−*Max*_ = 13

Regarding their rheological response, in Fig. 2.(b), we compare the elastic and viscous modulus of the final gel obtained for each mechanism at *ϕ*_*int*_ = 15%. The branched mechanism with a small elasticity modulus and slope equal to two, as well as a viscous modulus with slope equal to one, indicate a specific power-law behavior in the rheological response and has a liquid-like behavior. Thus, despite its network-like connectivity this mechanism is insufficient to model viscoelastic response. In contrast, the plateau in *G*′ observed for Low-Connected and Highly-Connected mechanisms evidence a final higher rigidity of the gel. In particular, the Highly-Connected mechanism exhibits a larger change in the elasticity modulus during the gelation process. Besides the highlighted differences among mechanisms, in their bond count and elasticity modulus, the differences in their bond distribution reveal disparities in the rate at which the gel consolidates. For example, when comparing the Highly-Connected and Low-Connected mechanisms in (see Fig. 2.(c)) with the same average number of bonds,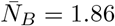, the Highly-Connected case exhibits faster gelation 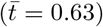 compared to the low-connected 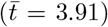 case. In this case, the larger number of bonds allowed during the crosslinking stage leads to a quicker consolidation or *fast gelation* of the Highly-Connected gel. Given that it depicts the solid phase in the Highly-Connected mechanism’s final state, we can anticipate that it will be able to replicate the pre-gelation, gel-point, and post-gelation phases of a gelation process.

For further analysis and calibration of fibrin-gel formation, we focus on the Highly-Connected mechanism due to its ability to accurately represent the entire gelation process, including a wide range of elasticity modulus variations during gelling, and faster gel network consolidation. While the Low-Connected mechanism can model the gelation process, it is better suited for slower gelation processes with lower elasticity. It’s important to highlight that the adopted gelling framework[26] supports modeling systems with features that span from low to high connectivity, by varying the degree of bond formation during the cross-linking stage. However, a detailed exploration of these parameters is beyond the scope of our current study. In subsequent sections, we address the identification of the three distinct gel phases —pre-gelation, gel-point, and post-gelation—using the Highly-Connected mechanism. Additionally, we compare the effect of gel concentration on the microstructural and rheological properties of the gel.

### 5.1 Gel phases identification

Identifying the gel point (GP) through viscoelastic characterization presents significant challenges. For accurate viscoelastic measurements, the sample must maintain a consistent degree of crosslinking or no mutational effects throughout the experiment. However, gelling systems are inherently transient, changing as they gel. To address this, experimental methods such as halting the crosslinking reaction before conducting rheological measurements,[39, 19] or employing multifrequency rheometric techniques, have been developed.[40, 41] Numerically, a key advantage in identifying the GP is our ability to separate the gelling process from rheological characterization. Instead of having to pause or stop the reaction to monitor the GP during gelling, we first run simulations up to the post-gelation stage. We save the particle positions of the system at specific time intervals. Then, as a post-processing step, we perform SAOS tests to observe changes in the tan(*δ*) at various frequencies for selected samples from the entire gelling trajectory. We perform a SAOS test on one selected sample. If the system is in pre-gel (primarily liquid-like behavior), move forward in time to a later snapshot with higher 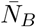(indicating more bonds and progression toward gelation). If the system is post-gel (solid-like behavior), move backward to an earlier snapshot with a lower 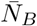. During the rheological characterization of these samples, we only consider existing hydrodynamic and bonding interactions, without allowing any new bond formations. This approach enables us to track changes in the system’s viscoelastic properties at different stages, free from the ongoing gelling process. We essentially confirm that no structural changes have occurred by asserting that the mutation number is zero.[19]

In Fig. 3.(a) we show tan(*δ*) for the three phases of gelation –pre-gelation, gel-point, and postgelation– at five different times (two at pre-gelation, two at post-gelation, and gel point) for the highest concentration, *ϕ*_*int*_ = 15%. In the pre-gelation phase, as the frequency (*ω*) increases, we observe a decrease in *tan*(*δ*), indicating behavior typical of a viscoelastic fluid. As gelation advances, the relationship between *δ* and *ω* weakens, and by the time we reach the gel point, *tan*(*δ*) becomes independent of frequency. Beyond the gel point, the material behaves as a viscoelastic solid, showing a characteristic dependency on frequency. In Fig. 3.(a), we additionally include the values of *D*_*f*_, 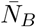 at the selected time sampled. Across the various systems evaluated, we identify that the average number of bonds between particles can be linked to the different phases of gelation. For instance, for *ϕ*_*int*_ = 15% a value of 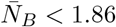 is indicative of the pre-gelation phase, while 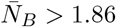 correlates with the post-gelation stages. In Fig. 3.(b), 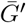 and 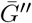 evolution in these phases are shown. Both increase with time but *G*′ shows a more noticeable increment, indicating the development of the gel network. The increase in *G*′ and *G*″ and the decrease in *D*_*f*_ after GP indicates microstructural changes after GP. The GP occurs at lower values of both 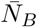 and *G*′ and these parameters continue to change well beyond the GP. We must note that for a fixed *ϕ*_*int*_, after the gel point the morphology of the network can keep changing due to cross-linking, without further increase of the gel volume fraction. This effect, leads to a morphological transition to more porous or open gels, along with a reduction in *D*_*f*_.

**Figure 3:**
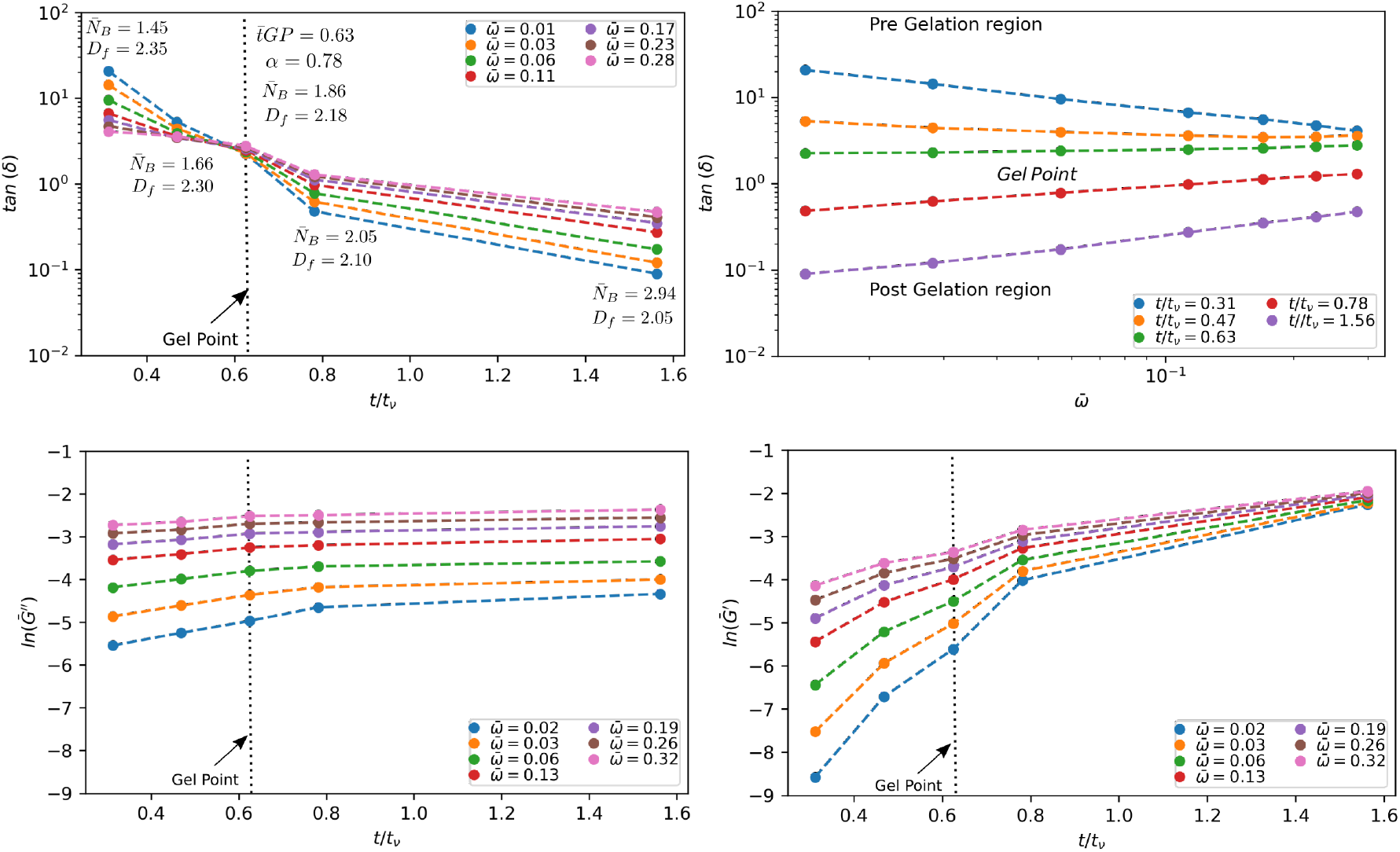
(a) *tan*(*δ*) at pre-gelation, gel-point, and post-gelation phases for *ϕ*_*int*_ = 15%, (b) 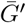 and 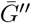evolution at pre-gelation, gel-point, and post-gelation phases for *ϕ*_*int*_ = 15%

### 5.2 Effect of thrombin concentration on gel microstructure and viscoelasticity

We focus now on the variation of the microstructure and viscoelastic properties of the gel, with the initial concentration of **P** or equivalently to the thrombin concentration during the formation of fibrin-thrombin gels.

#### 5.2.1 Microstructure behavior

Simulation snapshots of gel bond networks at GP are shown in Fig. 4.(a) for *ϕ*_*int*_ = 7.5, 10, 12.5, 15% gel concentrations. They show higher concentrations lead to more densely packed structures and less open. Higher thrombin concentrations have been shown to cause the formation of dense microstructure and fibers with a high degree of branching due to rapid polymerization. These structures are often described using terms such as “fine,” “coarse,” “open,” “sparse,” and “tight”. Although our model does not explicitly model fiber branching at the monomer level, we can qualitatively say that higher initial concentrations of passive particles result in a denser network with more branch points due to the greater number of particles.[13, 15] These effects on branching are consistent with SEM images of fibrin clots formed at different concentrations of thrombin, analyzed by Rsiman and coauthors, as shown in Fig. 4.(b).[15]

**Figure 4:**
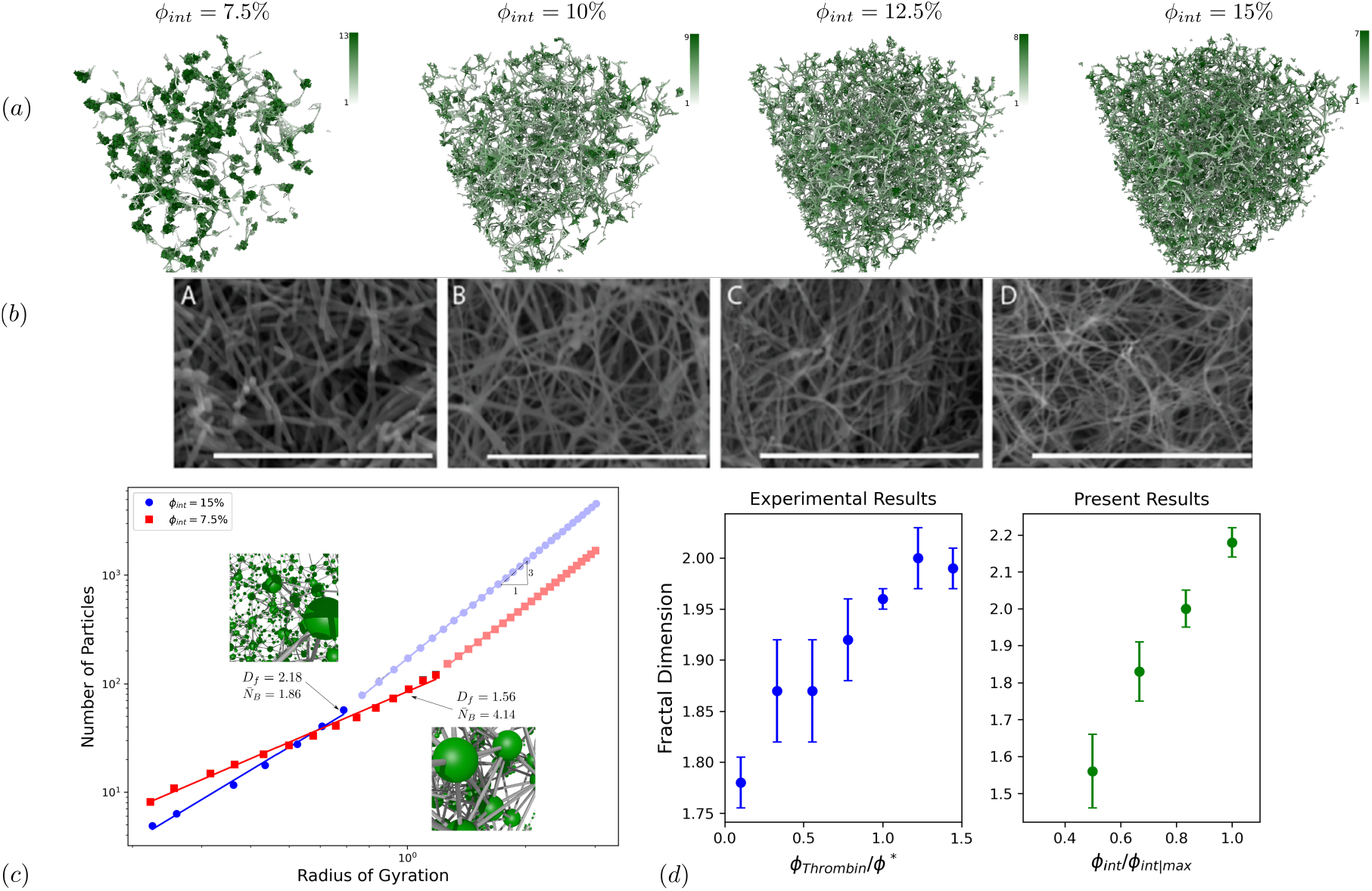
(a) Snapshots of gel bond networks at the GP for *ϕ*_*int*_ = 7.5, 10, 12.5, 15% concentrations. (b) SEM images of fibrin clots formed at different thrombin concentrations as imaged by Risman et al^1^[15] (c) Mass scaling of the clusters as a function of their radius of gyration (SDPD units). We obtain the fractal dimension from these data *D*_*f*_ = 1.56 *±* 0.1 for *ϕ*_*int*_ = 7.5% and *D*_*f*_ = 2.18 *±* 0.04 for *ϕ*_*int*_ = 15% (d) *D*_*f*_ as a function of concentration compared with experimental results.[11]

In general, a quantitative microstructure characterization is rarely performed.[8, 23] In most cases, the results of quantitative characterization are limited to the average distance between branch points and the densities of the branch points. Here we provide a quantitative analysis of the microstructure complexity and branching of the gels by measuring their fractal dimension *D*_*f*_. We compute *D*_*f*_ by considering the variation in the mass *M* of the formed clot with respect to their radius of gyration (*R*_*g*_) as 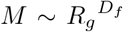 (see SI.S2.1 for a detailed description). The estimation of *D*_*f*_ is restricted on the length scales *R*_*g*_ *< ϵ*, where *ϵ* corresponds to the size of a blob in which *ϕ*_*blob*_ = *ϕ*_*int*_. Beyond this scale, the network is considered to be homogeneous, and *D*_*f*_ = 3.

In Fig. 4.(c) we present the measured *D*_*f*_ for the lowest and highest concentrations evaluated, indicating the range in *D*_*f*_ attained with our numerical model, where *D*_*f*_ = 1.56*±*0.1 for *ϕ*_*int*_ = 7.5% and *D*_*f*_ = 2.18*±* 0.04 for *ϕ*_*int*_ = 15%. At the lowest concentrations, we have low-mass gels with the largest number of bonds observed at the gel point, 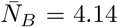. This is consistent with percolation models, network theory, and polymer physics, where increasing bond connectivity leads to more compact networks with lower fractal dimension.[42, 43]. As the particle concentration increases, *D*_*f*_ also rises because the gel occupies more space. Consequently, fewer bonds per particle are required to reach a percolating cluster in a densely connected network.

Within the range of *ϕ*_*int*_ investigated, we observe a quasi-linear proportionality of *D*_*f*_ with *ϕ*_*int*_. This is consistent with experimental observations[11] on the dependency of fractal dimension at the gel point with thrombin concentrations, as depicted in Fig. 4.(d). In Fig. 4.(d), we normalize both thrombin concentration (*ϕ*_thrombin_) and *ϕ*_*int*_ to allow a proper comparison between the experimental and numerical data. The thrombin concentration in the experimental data is normalized against the concentration where a plateau in the gel-time is reached,[11] corresponding to *ϕ*^∗^ = 0.09 NIH/ml. For simulation results, we use *ϕ*_*int*|*max*_ = 15% as a representative critical value. We must note that for larger particle concentrations –*ϕ*_*int*_ *>* 15%–, the simulated gels exhibit *D*_*F*_ *>* 2. Therefore, we select the threshold for the critical concentration to ensure consistency with experimental evidence for thrombin clots, where a fractal dimension much larger than 2 is unusual (See SI.S2 for further discussion about the selection of *ϕ*_*int*|*max*_ = 15%).This normalization provides a *ϕ*_*int*_ : *ϕ*_thrombin_ mapping, aligning the maximum gel concentration simulated, to the maximum effective thrombin concentration, beyond which the gelation kinetics is not affected.

#### 5.2.2 Viscoelastic behavior

In Fig. 5.(a) and (b), we summarize the viscoelastic characterization of the gels with varying concentrations. Values of the elastic *G*′ and loss modulus *G*″ at the gel point exhibit a power-law relationship with respect to frequency for all the concentrations evaluated, as depicted in Fig. 5.(a).

**Figure 5:**
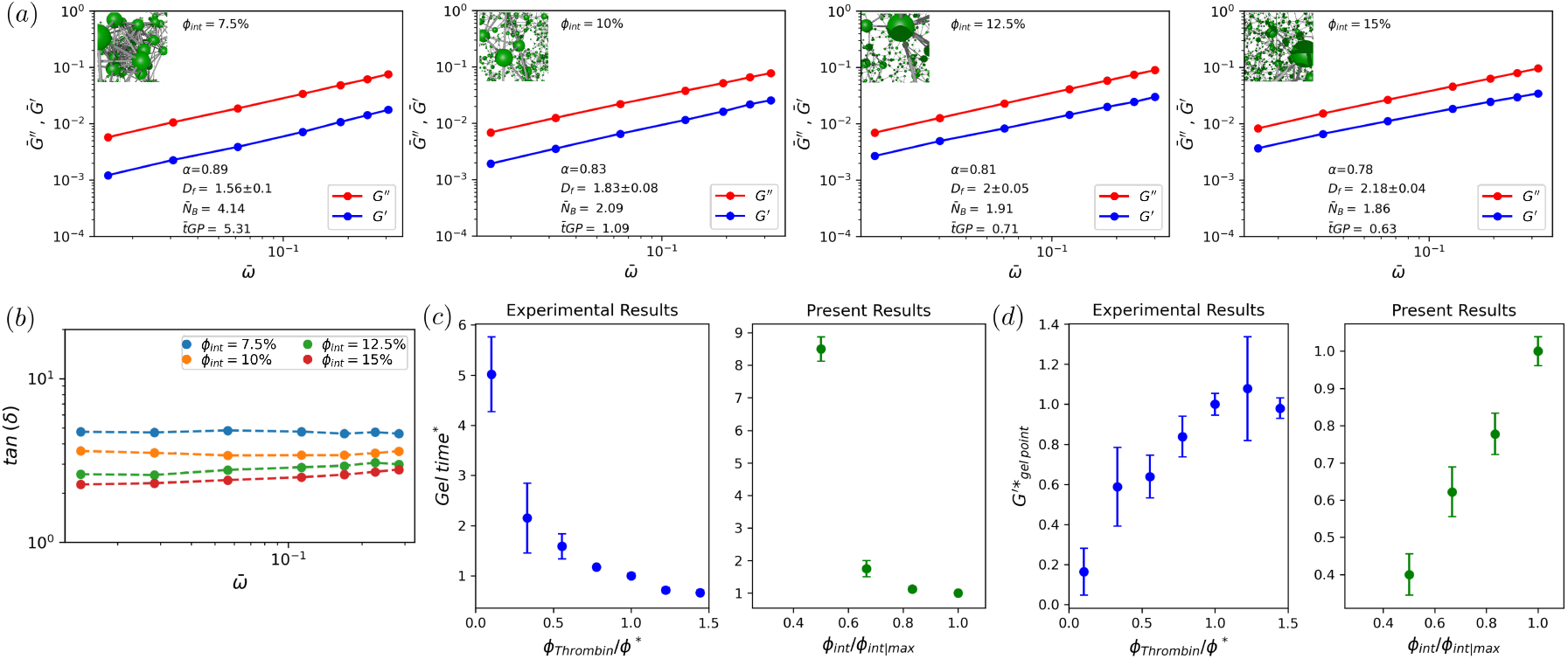
(a) Values of elastic modulus 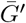, and loss modulus 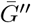 as a function of frequency for *ϕ*_*int*_ = 7.5, 10, 12.5, 15% concentrations. (b) *tan*(*δ*) = (*G*″*/G*′) *tan*(*απ/*2) as a function of frequency. (c)tGP as a function of concentration compared with experimental results. (d) 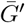 (measured at *ω* = 6.28*Hz*) at the GP as a function of concentration compared with experimental results (all values normalized based on their respective values observed at the *ϕ*_*int*|*max*_ = 15% for present results and *ϕ*^∗^ = 0.09 NIH/ml for experimental results.[11]

The corresponding values of *D*_*f*_, 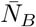, tGP, and *α* are also presented for comparison. In Fig. 5.(b), we show the variation of tan(*δ*) for the different concentrations, as a function of frequency. The loss tangent is frequency independent and approximately ∼ tan(*απ/*2),[19] indicating that our gels are approximately at the gel point. We can note that the small variations in *ϕ*_*int*_ (7.5% to 15%) cause significant changes in the microstructure, resulting in noticeable alterations in the viscoelastic modulus and gelation time. Both *G*′ and *G*″ increase with concentration, but the greater change in *G*′ leads to a reduction in tan(*δ*).

Overall, our simulations show that the changes in the viscoelastic properties of the gels, specifically the reduction in the power-law exponent and the rise in the storage modulus satisfactorily reproduce the response induced by thrombin in fibrin gels.[11]. In Fig. 5.(c) and (d), we further compare the values of tGP and 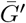(measured at *ω* = 6.28*Hz*) as a function of *ϕ*_*int*_, with previously reported evidence[11]. Following the normalization used rationalize the variation in *D*_*f*_, in Fig. 5.(c) and (d) the parameters tGP and *G*′ are normalized using the values gel-point time and elasticity modulus observed at the thrombin concentration, *ϕ*^∗^ = 0.09 NIH/ml, corresponding to the critical concentration where the gelation time reaches a plateau. For simulation results, we use *ϕ*_*int*|*max*_ = 15% as the critical value. In Fig. 5.(c), we identify that tGP quickly decreases as the concentration of thrombin increases, indicating faster gelation at higher concentrations. The variation in gelation rates—from slow at *ϕ*_*int*_ = 7.5% to fast at *ϕ*_*int*_ = 15%—is attributed to differences in the activation probability of the Highly-Connected mechanism, which speeds up the cross-linking process. As thrombin concentration increases, so does the rate of polymerization. We must note that similar to the experimental evidence, this reduction exhibits a limiting behavior, where further increments in thrombin concentrations do not affect the gelling time. The existence of this limiting value of tGP is attributed to the fact that in those experimental essays, thrombin concentration solely dictated the kinetics of fibrinogen activation. In this sense, the variation on *ϕ*_*int*_ in our model is equivalent to the variation in thrombin concentration in the experiments.

In Fig. 5.(d), we note that, contrary to the gelation time, the storage modulus increases with thrombin concentration. This trend suggests that gels formed at higher thrombin concentrations have a greater ability to withstand elastic stress, remaining within the linear viscoelastic region.

Experimentally, it has been observed that *G*′ plateaus, indicating that further increases in thrombin concentration do not affect the gel’s storage modulus beyond a critical concentration. This plateau is reached at the concentration of *ϕ*^∗^ = 0.09 NIH/ml, coinciding with the condition where gelation time also plateaus. Our model, which maps *ϕ*_*int*_ : *ϕ*_thrombin_, shows a satisfactory agreement with experimental variations of *G*′. For example, at a thrombin concentration of *ϕ*_*thrombin*_*/ϕ*^∗^ = 0.5, experiments show *G*′ at approximately 60% of its maximum, while simulations show it around 60%.

##### Relationship of power-law exponent *α* and fractal dimension

The relationship between the power-law exponent *α* and the fractal dimension *D*_*f*_, provides insights into the interplay between the viscoelastic properties and the microstructure of gels. This relationship offers a method to indirectly estimate the structural properties of gels through rheological characterization[2]. In Fig. 6, we present the variation of *D*_*f*_ with *α* for our simulated fibrin gels. Our results show *D*_*f*_ values ranging from 1.56 ± 0.1 to 2.18 ± 0.04 and *α* values from 0.89 to 0.78. The increase in *α* reflects a reduction in the degree of network branching within the gel, leading to an overall reduction in the fractal dimension of the gel. Higher *D*_*f*_ values indicate a more compact network, while lower *D*_*f*_ values suggest a more open network structure.

**Figure 6:**
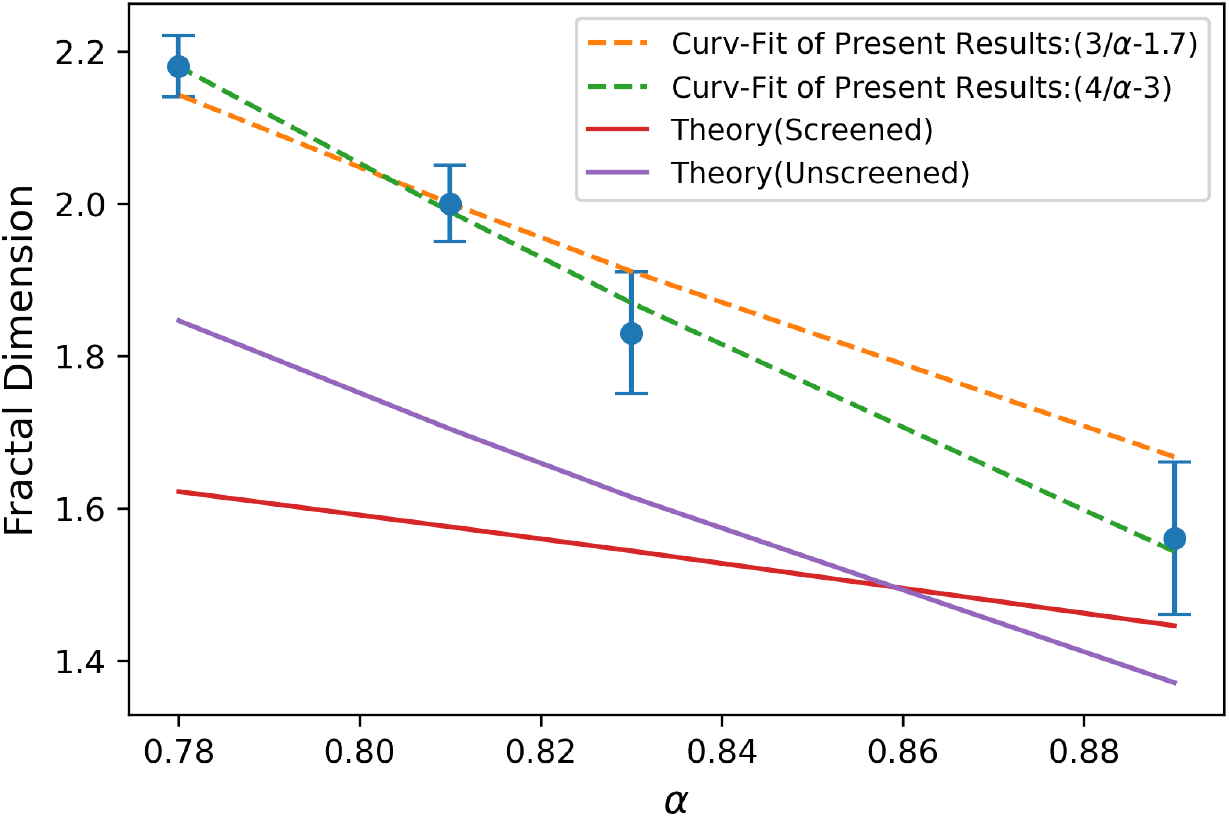
*D*_*f*_ as a function *α* in *d* = 3 for the unscreened, screened cases[44], and present results.

In the context of fibrin gels, the theoretical framework introduced by Muthukumar[44] for polymeric systems has been adopted to approximate the dependence of *D*_*f*_ with *α*.[2] According to Muthukumar, in conditions where hydrodynamic and excluded-volume interactions are screened, the relationship between *α* and *D*_*f*_ is given by *D*_*f*_ = (*d* + 2)(2*α* − *d*)*/*2(*α* − *d*), with *d* representing the Euclidean dimension of the space. For unscreened interactions, the model predicts a stronger decay in the fractal dimension, following the form *D*_*f*_ = *d/α* − 2. In Fig. 6, we include the theoretical predictions for both screened and unscreened conditions. In general, the theoretical model of Muthukumar predicts lower values of *D*_*f*_ for a given *α* in comparison to our numerical results. The origin of this discrepancy can stem from inherent features of the experimental or numerical techniques adopted.[45, 46, 47, 48] However, the functional dependence observed for the simulated fibrin gels suggests a consistent agreement with the unscreened model. A curve fitting of our simulations results shows a better adjustment to a linear dependence of the form *D*_*f*_ = (*d* + 1)*/α* − 3. We must note that within the clustering framework adopted, the SDPD method explicitly accounts for hydrodynamic interactions between all the interacting particles of the systems. Consequently, the formed gel is expected to resemble the unscreened type of dependency as observed in Fig. 6. Despite the discrepancies observed between the theoretical predictions and our numerical results, the identified correlations between *D*_*f*_ and *α* indicate that the interdependence between the viscoelastic properties and the microstructure of fibrin gels can be effectively captured within the framework of our model.

## Conclusion

This study introduces a mesoscopic model that simulates the formation of fibrin-thrombin gels, focusing on their rheological behavior and microstructural characteristics. Our model effectively replicates the complex fractal structure and viscoelastic properties observed in fibrin-thrombin gels at the gel point. One of the key aspects of our model is its ability to account for variations in the concentrations of thrombin into the structure of fibrin gels. The resulting gels have fibers with fewer branch points and larger pores at lower thrombin levels. Conversely, higher thrombin concentrations lead to denser fiber networks with more branch points and smaller pores. This variation in fiber structure directly influences the stiffness of the fibrin network, with higher thrombin levels generally leading to increased stiffness.

Furthermore, our model demonstrates that increasing the initial concentration of passive particles—akin to increasing thrombin levels—results in a decrease in the gelation time (tGP) and the power-law exponent (*α*), while the fractal dimension (*D*_*f*_) and elasticity modulus (*G*′) increase. These trends are consistent with both experimental and numerical data from previous studies, showing values of *D*_*f*_ ranging from 1.56 ± 0.1 to 2.18 ± 0.04. Notably, our results not only align qualitatively with existing evidence but also show good quantitative agreement. Additionally, our numerical model is able to capture the interplay between the viscoelastic properties and the microstructure of fibrin gels.

By accurately capturing the dynamics of fibrin polymerization and network formation, our model offers valuable insights for clinical and bioengineering applications requiring precise gelation control. Additionally, this mesoscale clot model sets the stage for future Lagrangian heterogeneous multiscale modelling[49] of clotting processes under physiological flow conditions.

## Author contributions

N.M and M.E conceived and supervised the project. E.Z designed the computational experiments and conducted the simulations and data processing. The original draft of the manuscript was written by E.Z and N.M. and all authors contributed to the final version of the manuscript. N.M. developed the numerical implementation. All the authors discussed and analyzed the results. All authors approved the final version of the manuscript.

## Conflicts of interest

There are no conflicts to declare.

## Data availability

The data supporting this article have been included in the main document and as part of the Supplementary Information.

## Acknowledgements

This research is supported by the Basque Business Development Agency through the “Mathematical Modeling Applied to Health” Project. Also by the Basque Government through the BERC 2022-2025 program and by the Ministry of Science and Innovation: BCAM Severo Ochoa accreditation CEX2021-001142-S / MICIN / AEI / 10.13039/501100011033.

## Supplementary Information

## S1 Supplementary Equations

The thermal fluctuation is included in the model by

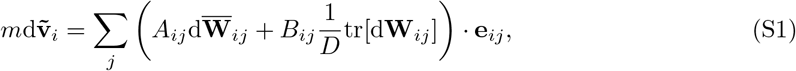

where 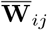 is a matrix of independent increments of a Wiener process for each pair *i, j* of particles, and 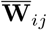 is its traceless symmetric part, given by

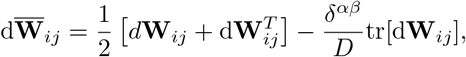

where *D* is the system dimensionality. To satisfy the fluctuation-dissipation balance the amplitude of the thermal noises *A*_*ij*_ and *B*_*ij*_ are related to the friction coefficients *a* and *b* through

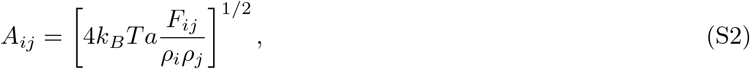

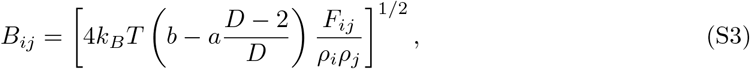

To describe the variation of the pressure with the density of the system we adopt the Cole equation (a.k.a. Tait’s equation of state) given by *p*_*i*_ = *c*^2^*ρ*_0_/7 [(*ρ*_*i*_*/ρ*_0_)^7^ 1] + *p*_*b*_ (S4) where *c* is the speed of sound on the fluid, and *ρ*_0_ is the reference density. The term *c*^2^*ρ*_0_*/*7 corresponds to the reference pressure of the system, given by 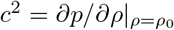. The parameter *p*_*b*_ is a background pressure that provides numerical stability by always keeping the system’s pressure positive.

We adopt the Lucy kernel[27] typically used in SDPD for the interpolant function,

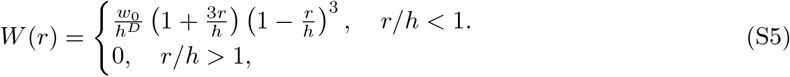

where *w*_0_ = 5*/π* or *w*_0_ = 105*/*16*π* for two or three dimensions, respectively. The reader is referred to[31] for a comprehensive description of the SDPD method.

## S2 Fractal dimension evaluation

Fractal geometry as a tool for assessing the intricacy of natural formations introduced by Mandelbrot.[50] When an item exhibits self-similarity across different length scales, denoted as scale-free behavior, it qualifies as a fractal. This characteristic is governed by power-law functions featuring a singular exponent, leading to a non-integer dimension termed *D*_*f*_. In cluster analysis, the computation of *D*_*f*_ from the radius of gyration (*R*_*g*_) involves a linear fit to *R*_*g*_ ∼ *M* ^*β*^ on a log-log scale. Here, *D*_*f*_ = 1*/β* is the estimated fractal dimension. We calculate the radius of gyration *R*_*g*_ of a cluster using 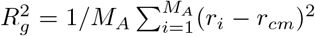, where *r*_*i*_ is the position of *i*th particle and *r*_*cm*_ is the center of mass of the cluster.

As particle volume fraction increases, the fractal dimension is expected to increase, approaching 3 when the volume fraction is 1.[51] Therefore, we limit our maximum concentration to *ϕ*_*int*_ = 15%, which gives us *D*_*f*_ *>* 2. Also, the fractal model can only be applied to particle volume fractions less than 20%. At higher volume fractions, the number of particles per cluster becomes so small that using a fractal model, and hence power law dependencies, is no longer justified.[51]

### S2.1 Fractal Dimension at Gel Point

Some studies[2] calculate the value of *D*_*f*_ from analysis of the viscoelastic data at the *GP* by the established relationship of Muthukumar for the incipient gel network (*D*_*f*_ = (10*α* −15)*/*(2*α* −6) for 3D space).[44] We use a numerical measure of the gel microstructure. The percolation theory defines the polymerizing system as macroscopically homogeneous on a length scale *L >> ϵ*. In contrast, for *L << ϵ* the sample-spanning network cluster is a ‘self-similar’ or fractal object. The blob-like network structure of gels has a profound impact on their features. A uniform distribution of particles is seen when the concentration of particles inside a blob is equal to the concentration of particles across the gel.

## S3 Simulation Details

We conducted numerical simulations using the clustering aggregation framework (based on the smoothed dissipative particle dynamics method SDPD) previously introduced.[26] We focus our investigations on periodic three-dimensional systems and consequently reported clusters with characteristic *D*_*f*_ ranging from 1.56 ± 0.1 −2.18 ± 0.04. Since the formed gel spans the whole simulation domain, which is periodic, we used an arbitrary boundary condition[52] approach for imposing the shear rate in SAOS (see S4). The characteristic parameters of the SDPD method are shown in the Table.S1. Simulations were performed using the software LAMMPS modified to incorporate the SDPD model.[53, 54] To account for the influence of the randomness of the bonding process, all the simulations are conducted over five realizations, initialized with different positions and velocities, in addition, we computed the standard deviation of all reported results.

## S4 Arbitary boundary conditions for imposing shear

We use the proposed boundary conditions by Moreno and Ellero.[52] This approach is suitable for conducting rheological studies where rheological characterization can be conducted using arbitrary flow types (simple shear, extensional, mixed flow) in periodic simulation domains. They adopt a domain decomposition scheme that considers the system made by a core region (where the stress is measured), a boundary condition region (where the characteristic type of flow is imposed), and buffer regions (that stabilize the system). We use buffer regions that expand over two-fold the size of the interpolation kernel to ensure the system’s stability and simulate up to a satisfactory agreement the properties of the fluid.[52] These boundary conditions are suitable for applying the oscillatory shear over the gels percolating the simulation domain. In the Fig. S1 sketch of the regions defined for SDPD simulations, the type of boundary conditions defined according to the target velocity field is shown. Parameters used for boundary conditions are shown in the Table.S2.

**Figure S1:**
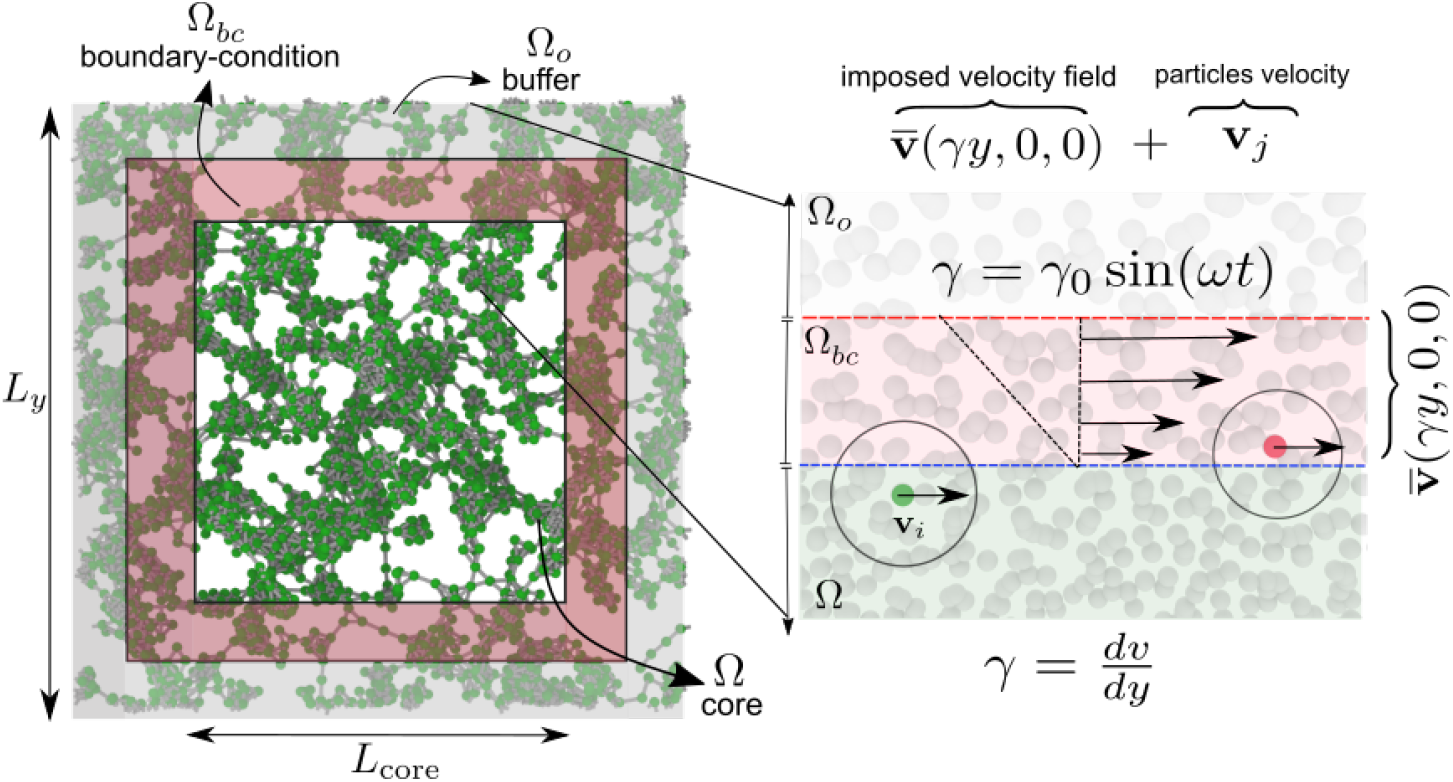
Sketch of the regions defined for SDPD simulations, and the type of boundary conditions defined according to the target velocity field. The interpolated velocity field acts over particles in Ω_*bc*_.[52]

## S5 Stress measurements and fluid inertia effect in oscillatory rheometry

We compute shear stresses on the core region (see arbitrary boundary conditions) of the simulation domain. With measured stress, we extract *G*″ and *G*′. For small-amplitude oscillatory shear, the stress response is also sinusoidal, but delayed by a phase delay, *δ* (*σ*(*t*) = *σ*_0_*sin*(*ωt* + *δ*)). We extract *G*″ and *G*′ by using the sine sum identity equation: *σ*(*t*) = *σ*_0_*sin*(*ωt*)*cos*(*δ*) + *σ*_0_*cos*(*ωt*)*sin*(*δ*). Then measured as *G*″ = *σ*_0_*/γ*_0_*sin*(*δ*) and *G*′ = *σ*_0_*/γ*_0_*cos*(*δ*). The total viscous stress measured *σ* includes contributions from both the solvent and the gel. However, to account only for the stress contributions associated with gel-gel and solvent-gel interactions, we subtract the viscous stress attributable to the solvent (*σ*(1 − *ϕ*_*gel*_)) from the total viscous part and keep the part of the gel. Here, *ϕ*_*gel*_ corresponds to the actual volume fraction of the gel (or active particles) at the time of stress measurement. We must note that in our simulations the control parameter is the initial concentration *ϕ*_*int*_, which in principle may not coincide with the final gel fraction, depending on the degree of conversion of passive to active particles. This difference occurs because, at the gel point, some **P** particles may remain.

It has been previously identified,[38] that spurious measurements in the stress at high frequencies and large gaps can emerge due to inertial effects. Since those effects are ubiquitous in numerical simulations, we subtract the apparent elastic modulus (*ρω*^2^*s*^2^*/*6) from the measured *G*′. This artificial term does not describe any rheological property and is composed of fluid density *ρ*, frequency *ω*, and gap width *s*, reflecting the influence of fluid inertia.[38]

## S6 Supplementary Tables

**Table S1:** Input parameters of the SDPD method.

|  |  |
| --- | --- |
| Domain size $[L_x \times L_y \times L_z]$ | $(40dx) \times (40dx) \times (40dx)$ |
| Total number of particles<br>($N_t$) | 64000 |
| Mass($m$) | 0.04 |
| viscosity($\eta$) | 10 |
| $k_B T$ | 0.000001 |
| Density($\rho_0$) | 1 |
| Pressure($p_0$) | 0 |
| Speed of sound ($c_s$) | 50 |
| Time step ($dt$) | $10^{-4}$ |
| Initial lattice spacing($dx$) | 0.2 |
| Cutoff radius ($h$) | $4dx$ |

**Table S2:** Parameters of arbitrary boundary condition.

|  |  |
| --- | --- |
| $L_{core}$ | $(20dx)$ |
| $L_y$ | $(40dx)$ |

## Footnotes

1 Reprinted from Risman et al., “Comprehensive Analysis of the Role of Fibrinogen and Thrombin in Clot Formation and Structure for Plasma and Purified Fibrinogen,” *Journal of Biomolecules*, 2024,14,230, DOI: 10.3390/biom14020230, licensed under CC BY 4.0. (https://creativecommons.org/licenses/by/4.0/)

## Notes

### Competing Interest Statement

The authors have declared no competing interest.

## References

[1] J. W. Weisel, Journal of Thrombosis and Haemostasis, 2007, 5, 116–124.

[2] P. A. Evans, K. Hawkins, R. H. Morris, N. Thirumalai, R. Munro, L. Wakeman, M. J. Lawrence and P. R. Williams, Blood, The Journal of the American Society of Hematology, 2010, 116, 3341–3346.

[3] D. Curtis, M. Brown, K. Hawkins, P. Evans, M. Lawrence, P. Rees and P. Williams, Journal of non-newtonian fluid mechanics, 2011, 166, 932–938.

[4] S. N. Stanford, A. Sabra, L. D’Silva, M. Lawrence, R. H. Morris, S. Storton, M. R. Brown, V. Evans, K. Hawkins, P. R. Williams et al., BMC neurology, 2015, 15, 1–8.

[5] J. Collet, D. Park, C. Lesty, J. Soria, C. Soria, G. Montalescot and J. Weisel, Arteriosclerosis, thrombosis, and vascular biology, 2000, 20, 1354–1361.

[6] M. Wygrecka, A. Birnhuber, B. Seeliger, L. Michalick, O. Pak, A.-S. Schultz, F. Schramm, M. Zacharias, G. Gorkiewicz, S. David et al., Blood advances, 2022, 6, 1074–1087.

[7] P. Evans, K. Hawkins, M. Lawrence and P. Williams, BIORHEOLOGY, 2008, pp. 130–130.

[8] D. J. Curtis, P. R. Williams, N. Badiei, A. I. Campbell, K. Hawkins, P. A. Evans and M. R. Brown, Soft Matter, 2013, 9, 4883–4889.

[9] N. Badiei, A. Sowedan, D. Curtis, M. Brown, M. Lawrence, A. Campbell, A. Sabra, P. Evans, J. Weisel, I. Chernysh et al., Clinical Hemorheology and Microcirculation, 2015, 60, 451–464.

[10] S. L. Rowe, S. Lee and J. P. Stegemann, Acta biomaterialia, 2007, 3, 59–67.

[11] D. J. Curtis, PhD thesis, Swansea University, 2012.

[12] R. I. Litvinov and J. W. Weisel, Matrix Biology, 2017, 60, 110–123.

[13] A. S. Wolberg, Blood reviews, 2007, 21, 131–142.

[14] A. Sabra, M. J. Lawrence, D. Curtis, K. Hawkins, P. R. Williams and P. A. Evans, Clinical Hemorheology and Microcirculation, 2020, 74, 147–153.

[15] R. A. Risman, H. A. Belcher, R. K. Ramanujam, J. W. Weisel, N. E. Hudson and V. Tutwiler, Biomolecules, 2024, 14, 230.

[16] A. Takahashi, R. Kita, T. Shinozaki, K. Kubota and M. Kaibara, Colloid and Polymer Science, 2003, 281, 832–838.

[17] R. Bateman, H. Leong, T. Podor, K. Hodgson, T. Kareco and K. Walley, Microscopy and Microanalysis, 2005, 11, 1018–1019.

[18] M. Brown, D. Curtis, P. Rees, H. Summers, K. Hawkins, P. Evans and P. Williams, Chaos, Solitons & Fractals, 2012, 45, 1025–1032.

[19] H. H. Winter and F. Chambon, Journal of rheology, 1986, 30, 367–382.

[20] P. A. Evans, M. Lawrence, R. K. Morris, N. Thirumalai, R. Munro, L. Wakeman, A. Beddel, P. R. Williams, M. Barrow, D. Curtis et al., Rheologica acta, 2010, 49, 901–908.

[21] A. Zakharov, M. Awan, A. Gopinath, S.-J. J. Lee, A. K. Ramasubramanian and K. Dasbiswas, Science Advances, 2024, 10, eadh1265.

[22] N. Moreno, J. E. Perilla, C. M. Colina and M. Lísal, Molecular Physics, 2015, 113, 898–909.

[23] S. Yesudasan, X. Wang and R. D. Averett, Journal of molecular modeling, 2018, 24, 1–14.

[24] N. Moreno, S. P. Nunes and V. M. Calo, Computer Physics Communications, 2015, 196, 255– 266.

[25] S. Yesudasan, X. Wang and R. D. Averett, Biomechanics and Modeling in Mechanobiology, 2018, 17, 1389–1403.

[26] E. Zohravi, N. Moreno and M. Ellero, Soft Matter, 2023, 19, 7399–7411.

[27] P. Espanol and M. Revenga, Physical Review E, 2003, 67, 026705.

[28] P. M. Morse, Physical review, 1929, 34, 57.

[29] T. C. Day, P. Márquez-Zacarías, P. Bravo, A. R. Pokhrel, K. A. MacGillivray, W. C. Ratcliff and P. J. Yunker, Biophysics reviews, 2022, 3, 021305.

[30] M. H. Periayah, A. S. Halim and A. Z. M. Saad, International journal of hematology-oncology and stem cell research, 2017, 11, 319.

[31] M. Ellero and P. Español, Applied Mathematics and Mechanics, 2018, 39, 103–124.

[32] N. Moreno, P. Vignal, J. Li and V. M. Calo, Procedia Computer Science, 2013, 18, 2565–2574.

[33] D. A. Fedosov, M. Peltomäki and G. Gompper, Soft Matter, 2014, 10, 4258–4267.

[34] D. N. Simavilla and M. Ellero, Journal of Non-Newtonian Fluid Mechanics, 2022, 305, 104811.

[35] R. B. Bird, R. C. Armstrong and O. Hassager, 1987.

[36] C. W. Macosko, Measurements and Applications, 1994.

[37] A. Vázquez-Quesada, M. Ellero and P. Espanol, Microfluidics and nanofluidics, 2012, 13, 249– 260.

[38] G. Böhme and M. Stenger, Journal of Rheology, 1990, 34, 415–424.

[39] F. Chambon and H. H. Winter, Polymer Bulletin, 1985, 13, 499–503.

[40] M. Geri, B. Keshavarz, T. Divoux, C. Clasen, D. J. Curtis and G. H. McKinley, Physical Review X, 2018, 8, 041042.

[41] R. E. Hudson-Kershaw, M. Das, G. H. McKinley and D. J. Curtis, Journal of Non-Newtonian Fluid Mechanics, 2024, 105307.

[42] P.-G. De Gennes, Scaling concepts in polymer physics, Cornell university press, 1979.

[43] D. Stauffer and A. Aharony, Introduction to percolation theory, Taylor & Francis, 2018.

[44] M. Muthukumar, Macromolecules, 1989, 22, 4656–4658.

[45] P. Meakin, Annual Review of Physical Chemistry, 1988, 39, 237–267.

[46] T. Hagiwara, H. Kumagai and K. Nakamura, Food Hydrocolloids, 1998, 12, 29–36.

[47] S. S. Narine and A. G. Marangoni, Food research international, 1999, 32, 227–248.

[48] L. de Martín, A. Fabre and J. R. Van Ommen, Chemical Engineering Science, 2014, 112, 79–86.

[49] N. Moreno and M. Ellero, Journal of Fluid Mechanics, 2023, 969, A2.

[50] B. B. Mandelbrot, New York, 1983.

[51] H. Wu and M. Morbidelli, Langmuir, 2001, 17, 1030–1036.

[52] N. Moreno and M. Ellero, Physics of Fluids, 2021, 33, 012006.

[53] S. Plimpton, Journal of computational physics, 1995, 117, 1–19.

[54] M. Jalalvand, M. A. Charsooghi and S. Mohammadinejad, Computer Physics Communications, 2020, 255, 107261.

